# CropStabDB: a protein stability resource and Engineering Atlas of orthologue-supported substitutions for 30 plant proteomes

**DOI:** 10.64898/2026.10.01.756114

**Authors:** Md. Isrfil Hossen, Yilin Pan, Ping Lan

**Author notes:** Corresponding author: Ping Lan. Mailing address: Institute of Soil Science, Chinese Academy of Sciences, No.298 Chuangyou Road, Nanjing, Jiangsu, 211135, P.R. China.

## Abstract

Protein stability, the capacity to maintain a folded structure and function, matters as much for transporters and regulators as for enzymes, and heat erodes it. Wheat, rice, maize and soybean supply two-thirds of human caloric intake, and warming reduces yields. Rubisco activase variants selected for thermostability increased photosynthesis, biomass and yield under heat. They came from gene shuffling; no method proposes residues at proteome scale. Genome editing has made such changes routine, yet predictors estimate melting temperature, name no residue, and have not been reported for a plant proteome. Here we present CropStabDB: 1,425,716 proteins from 30 plant species scored on six interpretable components, with an Engineering Atlas of 2,097,576 candidate substitutions in 51,440 proteins, 785,622 at Engineering Confidence Score ≥ 0.70. Every candidate is a residue a more stable orthologue carries at that column. Substitution-type scores match the direction of measured folding free energies in 842 of 926 domains (90.9%). On three proteomes from families absent from training, the model reaches an area under the curve of 0.919. Applied blind to wheat, the score reproduces the measured thermostability ranking of the Rubisco activase isoforms in all three subgenomes. CropStabDB is freely available without registration at https://cropstab.org.

## Introduction

Protein stability (whether a polypeptide adopts and maintains a folded conformation) is a precondition for function in every class of protein [1]. Enzymes are the familiar case, but transporters, receptors, structural proteins and transcriptional regulators lose activity on the same terms. Heat is what breaches it: every protein has a temperature above which it loses its fold and, with it, its activity. In crops the consequences reach field scale. Wheat, rice, maize and soybean supply two-thirds of human caloric intake, and each degree of warming is estimated to reduce their yields by 6.0%, 3.2%, 7.4% and 3.1% respectively [2]. The dependence is not confined to these four crops: it holds across the cereals, grain legumes, oilseeds, fibre, root, vegetable and fruit species of world agriculture.

Rubisco activase shows what this means for a single protein: it is progressively inhibited above the photosynthetic optimum, and the resulting fall in Rubisco activation state accounts for the temperature response of net photosynthesis [3,4]. In *Arabidopsis*, lines expressing activase variants selected for improved thermostability showed higher photosynthetic rates, higher biomass and increased seed yields than lines expressing the wild-type activase under moderate heat [5]. Those variants came from gene shuffling, a combinatorial recombination of related sequences followed by selection [6], not from any prediction of which residue to change. The residue to change can instead be read from a more stable relative: eleven substitutions taken from an alignment of the thermostable wheat and rice activase isoforms against warm-adapted species raised a heat-labile wheat isoform’s thermal midpoint by more than 7 °C without loss of catalytic velocity [7]. That alignment was built by hand, for one protein. Genome editing makes such substitutions routine in rice, wheat and maize [8], and the limiting step is the choice of residue: a typical protein admits thousands of possible substitutions, and an experiment can evaluate a few.

Sequence-based stability predictors address a different question. ProTstab2 [9], DeepTM [10] and DeepSTABp [11] estimate melting temperature, a bulk property of the folded state. Their reported applications are to animal proteomes and to thermophilic and industrial enzymes, not to plants. A melting temperature identifies no residue, and says nothing about which part of the sequence produced it. Repeating that alignment by hand for every protein is impractical, but orthologue sets spanning dozens of sequenced plant genomes can now be built automatically, so the comparison that yielded eleven substitutions for one protein can be made for every protein at once. Two further developments make the result testable at that scale. The AlphaFold Protein Structure Database [12] provides per-residue model-confidence estimates for proteomes across the tree of life, a proteome-scale record of how confidently each region is predicted. Mega-scale proteolysis has measured folding free energies for around 776,000 variants of natural and designed protein domains 40–72 residues in length [13], an experimental standard against which a predicted substitution effect can be tested.

Here we present CropStabDB, which scores 1,425,716 proteins from 30 plant species in 12 families on six interpretable components derived from 64 sequence features, six numbers for every protein rather than one (**Figure 1**). Its Engineering Atlas reports 2,097,576 candidate substitutions across 51,440 proteins (**Figure 2**). Each candidate is a residue carried by a more stable orthologue at the aligned column, restricting proposals to substitutions already tolerated in a living plant rather than generated *de novo*. Applied blind, the score reproduces the measured thermostability ranking of the wheat Rubisco activase isoforms. CropStabDB is freely available without registration at https://cropstab.org, and is deposited with a permanent identifier.

**Figure 1.**
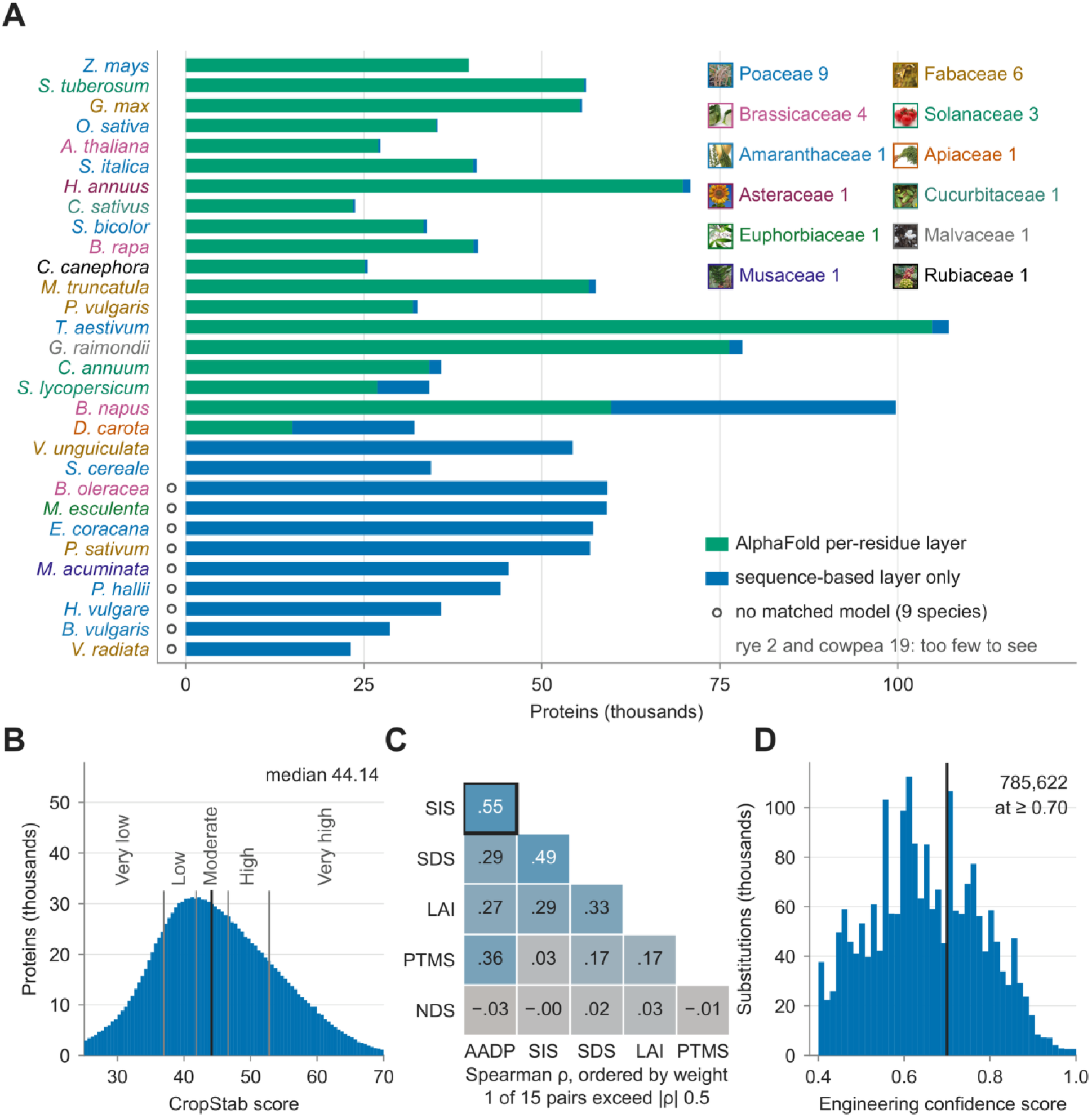
What the resource contains · **A** proteins per species, split into the AlphaFold-layer and sequence-only shares, species labelled by family with the twelve family photographs as the legend, and an open marker on the nine species with no matched model at all, distinct from rye (2 proteins) and cowpea (19), whose AlphaFold shares are too small to see · **B** the CropStab score across all 1,425,716 proteins, with the four quintile cut points and the median; lower values indicate the more stable direction · **C** Spearman correlation between the six components · **D** the Engineering Confidence Score across all 2,097,576 candidate substitutions, with the 0.70 threshold

**Figure 2.**
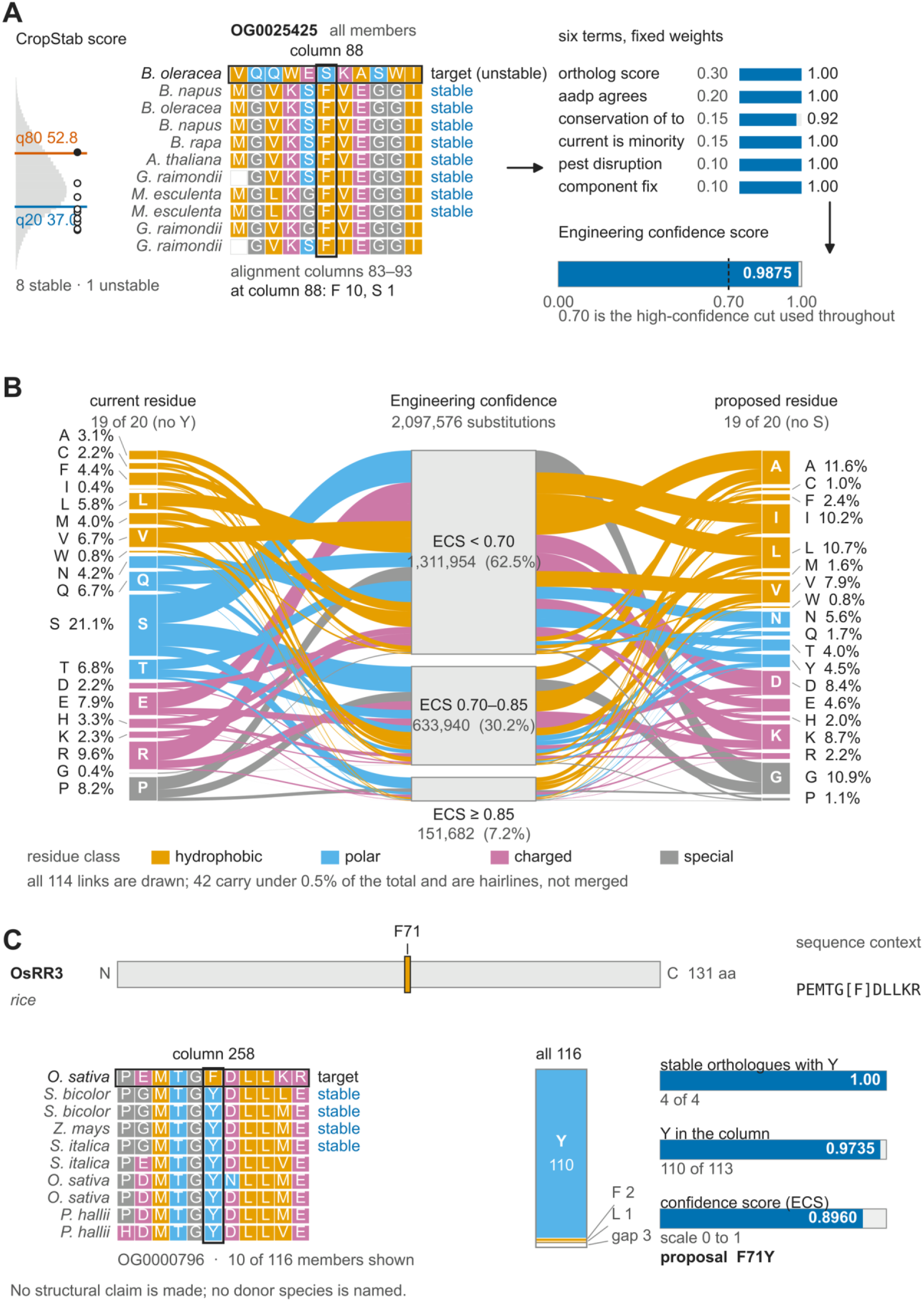
The Engineering Atlas · **A** mechanism, on a real alignment · **B** residue → confidence band → residue · **C** OsRR3 F71Y on its alignment column

## Database construction and content

### Sequence data and filtering

Every value in CropStabDB is computed from sequence alone. The 30 proteomes were obtained on 24 March 2026 as complete protein FASTA files, 29 from Ensembl Plants [14] and one from NCBI; **Table 1** lists each species, family, assembly and download source. The files are not deposited, because the providers’ terms differ, but their names and checksums are (**Table S1**). Sequences were uppercased, asterisks and ambiguity characters removed and residues outside the 20 standard amino acids discarded; entries left with fewer than 30 residues were dropped. Of 1,428,278 input sequences, 1,425,716 (99.8%) passed and form the database (**Figure S1**). Features and stored lengths refer to the cleaned sequence; 3,687 proteins lost one or more characters, and the script regenerating the cleaned form from the provider files is deposited.

**Table 1.**
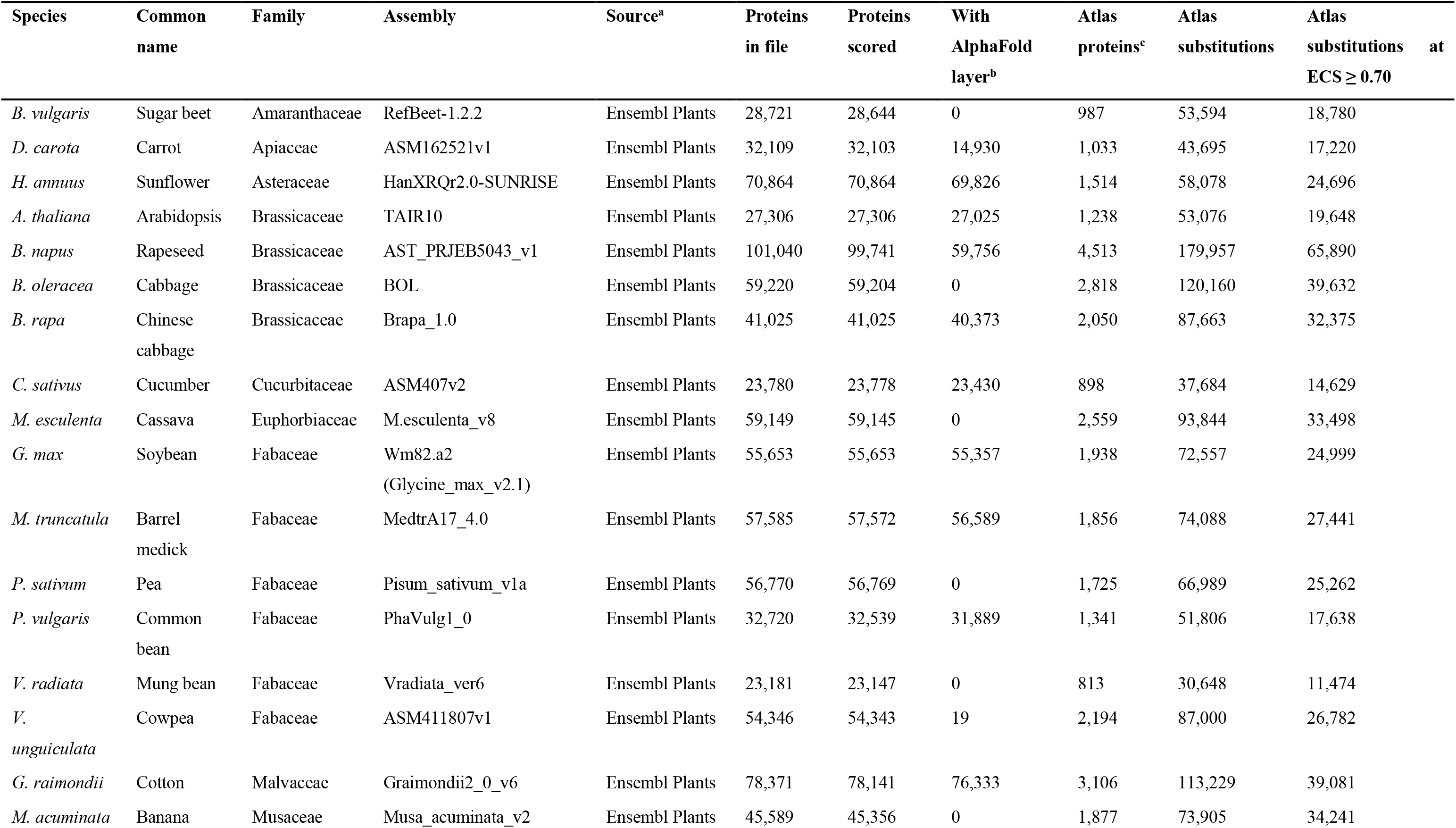
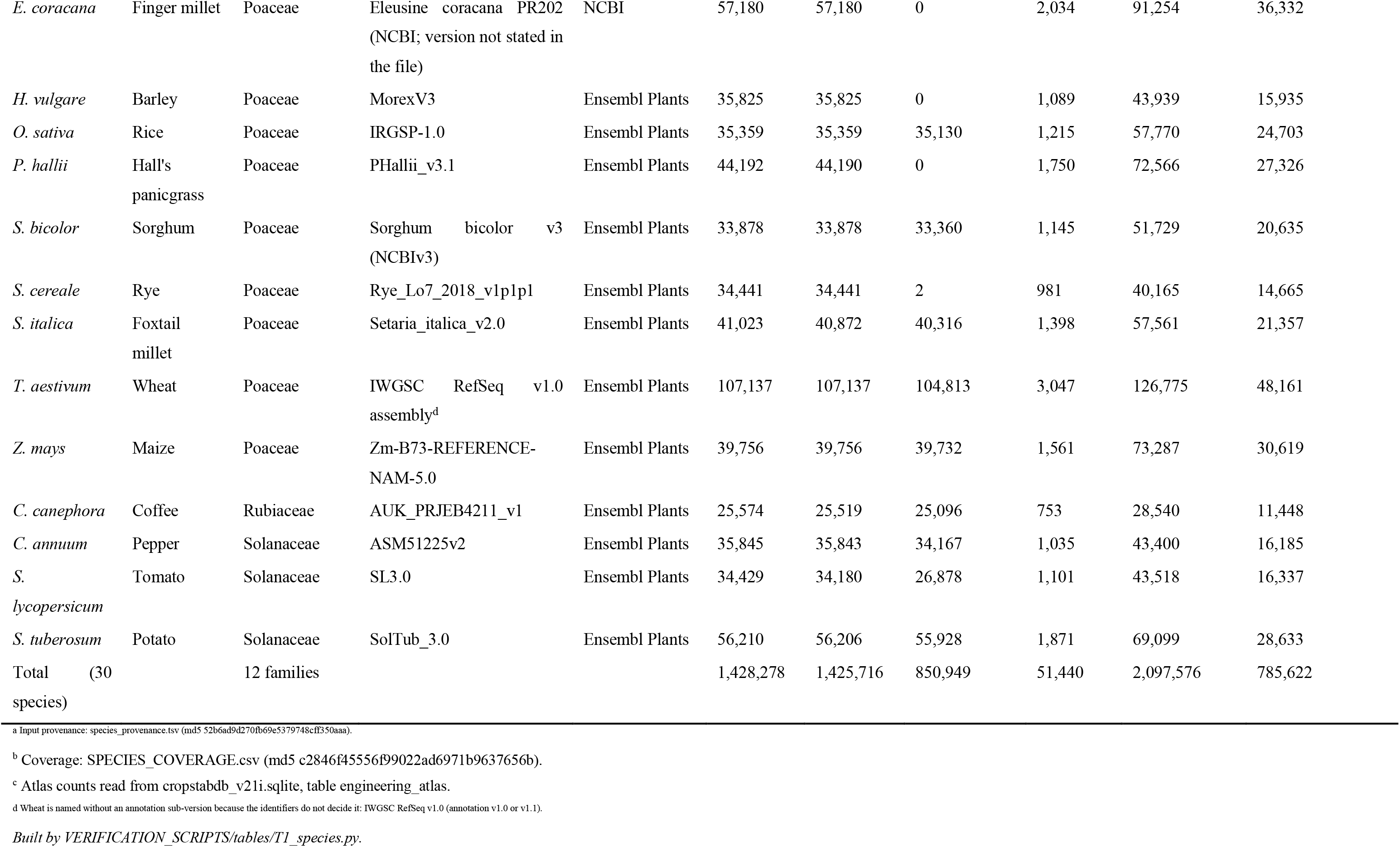
Species, assembly, protein counts and coverage The 30 species of CropStabDB, with the assembly named in the input file’s own headers, the number of protein records in that file, the number scored after filtering, the number carrying the AlphaFold per-residue confidence layer, and the number of proteins and candidate substitutions each contributes to the Engineering Atlas.

### Sequence features and component scores

The score is the weighted sum of six components, each computed from a defined subset of the features and interpretable on its own. We computed physicochemical, compositional and motif-based sequence features and reduced them in two steps, 74 to 71 to 64; the candidate list, intermediate lists and group ablation record are in **Table S2**. The full feature dictionary is deposited. The six are: a sequence-intrinsic score (SIS), a structural-disorder score (SDS), a post-translational-modification score (PTMS), an N-terminal descriptor score (NDS), an amino-acid-disposition profile (AADP) and a predicted-compartment weight, stored under the column name LAI. SIS is a logistic scaling of a dipeptide-composition index [15,16]; SDS rescales the mean TOP-IDP disorder propensity [17] over a 21-residue sliding window onto 0–1; PTMS is the rank percentile of the density of S, T, Y and K residues and N-X-S/T sequons. The 20 AADP residue weights come from SHapley Additive exPlanations (SHAP) attributions [18] of the released model over a sample of proteins, as w = mean|SHAP| × sign(r), where r is the correlation between that residue’s frequency and the model output; the weights are deposited as a parameter file. The NDS values are listed in **Table S3** and implemented in the released scoring script. The six components combine into the CropStab score on a 0–100 scale, lower values indicating a greater predicted propensity to adopt and maintain a confidently folded conformation; the proteome-wide median is 44.14.

### Classifier

One reference target sets both the classifier’s label and the component weights: AlphaFold’s own per-residue confidence [12]. The released classifier is a random forest of 200 trees (max_features=’sqrt’, class_weight=’balanced’, random_state=42); no hyperparameter search was performed. It was trained on 851,077 proteins from the 21 species with matched AlphaFold models, the label being a mean predicted local distance difference test (pLDDT) score below 70. Per-residue pLDDT separates disordered regions from ordered ones [19], making the threshold a reasonable label. The label is a model-confidence threshold, not a thermodynamic measurement, so the score is a predicted propensity to be confidently folded; its relation to measured folding free energy is established directionally below. Two of those species contribute almost nothing: cowpea has 19 labelled proteins and rye 2, so the leave-one-species-out evaluation covers 19 species rather than 21. A per-residue layer is stored for 850,949 of the 851,077; the remaining 128 are *Daucus carota* proteins carrying a training label whose per-residue columns the database did not retain. Software versions are pinned in the deposit, and the serialised model reproduces its stored predictions across all 1,425,716 proteins.

The component weights were set proportional to each component’s discriminative skill against the same AlphaFold-derived target the model is trained on, taken as |2 × AUC − 1| over 851,077 proteins, where AUC is the area under the receiver operating characteristic (ROC) curve; component correlations are in **Table S4**, values and weights in **Table S5**. The combined score reaches AUC 0.7316 against that same target. Because the weights were fitted to it, that figure is an internal consistency check and is reported here rather than among the validation results. Because skill was taken as an absolute value, NDS received a positive weight of 0.03 (AUC 0.4766 against that target, below chance) despite separating the classes in the wrong direction. It is therefore reported as a descriptive component of N-terminal composition and never as a discriminative one; setting its weight to zero changes the score’s AUC by +0.0004 and leaves protein ranking effectively unchanged (Spearman 0.9996).

### Structural confidence reference layer

For 850,949 of those proteins the model’s per-residue confidence was retained and is stored as data; only the protein-level mean entered the classifier, as the training label above. Models were linked by UniProt accession [20], Ensembl identifier [14] or gene name, recording the fraction of residues below pLDDT 50, 70 and 90, the longest run below 70, the number of low-confidence regions of at least 30 residues, and the mean pLDDT. The remaining 574,767 proteins carry a sequence-based disorder layer, and every protein records which it has. Coverage is uneven rather than partitioned by species: nine of the 30 have no matched models, and in the other 21 gaps range from a few dozen proteins to most of a proteome (**Figure 1A**). Coverage therefore does not bias the score: the per-species median is similar in those nine species and the other 21 (**Figure S2**).

### Identifying candidate substitutions

A candidate substitution is a residue that a more stable orthologue of the same protein carries at the aligned column. Orthogroups were inferred with OrthoFinder 3.1.2 [21] (DIAMOND search [22], FAMSA alignment [23], MCL inflation 1.2), and its trimmed alignments used to find those residues at every position of every protein (**Figure 2A**). Stable and unstable members are set by two fixed thresholds, the 20th and 80th percentiles of the score across all 1,425,716 proteins (36.99 and 52.81); within an orthogroup, members below the lower threshold are stable and those above the upper are unstable. Identifiers were reconciled through a crosswalk table, not by string matching, because conventions differ between sources. Each candidate is scored by an Engineering Confidence Score (ECS), a fixed-weight sum of six terms: the share of the orthogroup’s stable members carrying the proposed residue (0.30), agreement with the amino-acid-disposition expectation (0.20), the proposed residue’s frequency in the alignment column (0.15), whether the current residue is itself a minority in that column (0.15), disruption of a PEST motif, a segment rich in proline, glutamate, serine and threonine (0.10), and a coarser threshold on the same amino-acid-disposition difference (0.10), so that six terms are formed from five distinct inputs. All six are defined in **Table S6** and implemented in the deposited script. Each term rises monotonically across ECS bands (**Figure S3**).

Alignment columns were converted to residue positions and verified against the alignments for every row; rows whose protein did not map to the source FASTA (230) or whose mapped length disagreed with the stored length (150) were removed. Each entry carries an 11-residue context string with the focal residue bracketed, so a position can be located without counting, and a flag marking the 12,705 whose context is not unique within their protein. The score describes an isolated sequence, so it does not account for chaperone assistance, cofactor or metal binding, the actual modification state of a residue, or stabilisation within a complex; a substitution that disrupts a binding site can be destabilising in the cell while leaving the score unchanged. Every Atlas column is defined in **Table S6**.

### Scores and classes

CropStabDB scores all 1,425,716 proteins and places each in one of five classes bounded by quintiles of that distribution (**Figure 1B**), so class membership is comparable across species even though the proteomes differ in size and composition. Every record also carries the six component values and the 64 features they come from. Across all proteins only one of the fifteen component pairs exceeds |ρ| = 0.5 (**Figure 1C**). A user therefore sees not only which class a protein falls in but which components put it there. As a check on how the score behaves, most of its variance sits between orthogroups rather than species: 74.9% against 1.0%, from two one-way decompositions over the 1,156,886 representative isoforms in the 60,214 orthogroups with more than one member. A protein’s score therefore reflects its family more than its species, and orthologues score similarly across species (ρ = 0.7244 over 45,043 pairs, against +0.0077 for shuffled pairs). These and the other quality-control figures quoted in the text are collected, with their n and script, in **Table S7**.

### The Engineering Atlas

The Atlas turns the score into a decision: which residue to change. It contains 2,097,576 candidate substitutions across 51,440 proteins, of which 785,622 (37.5%) reach the high-confidence threshold of ECS ≥ 0.70 (**Figure 1D**, **Figure 2B**). They fall in 6,223 orthogroups: those of the 60,214 with more than one member that hold both a stable and an unstable member. The Atlas is sparse: of the 380 possible ordered substitution types only 208 are ever proposed (**Figure S4**), the largest single cell serine to alanine with 54,984 entries, a consequence of the orthologue constraint.

One entry illustrates the output (**Figure 2C**). In the rice type-A response regulator OsRR3 [24], the Atlas proposes F71Y at residue 71 of 131, with the context string PEMTG[F]DLLKR. Every stable orthologue carries tyrosine at the aligned column (fraction 1.00), tyrosine accounts for 97.3% of that column overall, and the Engineering Confidence Score is 0.896. The position lies in the protein’s interior, not at either terminus, so the substitution is not a trivial consequence of terminal variability.

### Validation

Validation is reported at three levels of independence (**Figure 3**).

**Figure 3.**
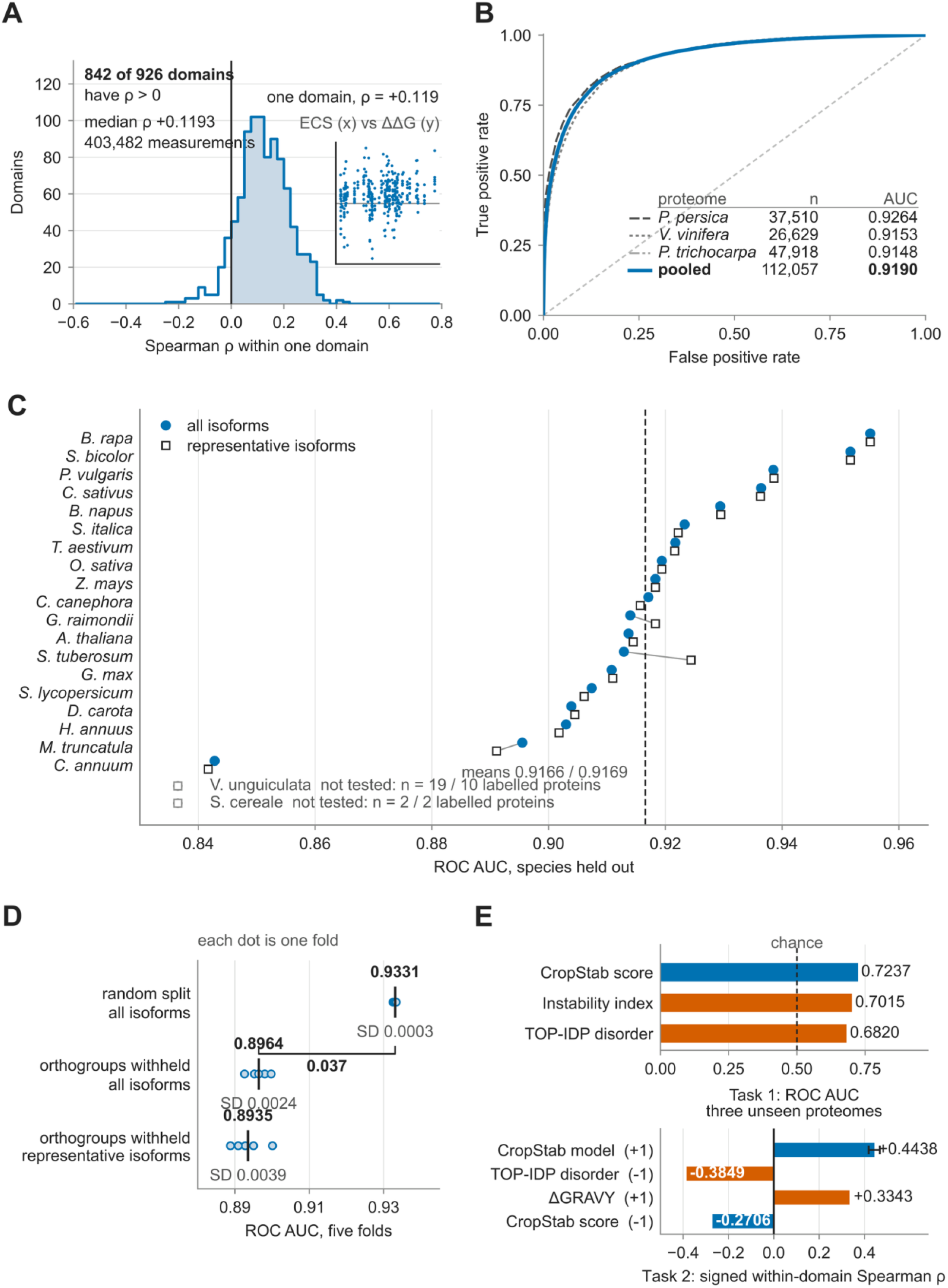
Validation and benchmark · **A** measured ΔΔG, positive values stabilising · **B** three unseen proteomes · **C** leave-one-species-out · **D** cross-validation folds · **E** the benchmark, both tasks

#### Measured folding free energies

Substitution-type mean ECS values were compared against measured ΔΔG from mega-scale proteolysis experiments on small protein domains [13], on the distributed convention that ΔΔG = ΔG(mutant) − ΔG(wild type) and positive values are stabilising. Matching is on residue identity only; position is never used, so the comparison is independent of the resource’s own residue numbering. Across 926 domains with at least 20 matched measurements and three distinct ECS values, 842 (90.9%) show a correlation in the declared direction (binomial P = 1.91 × 10⁻¹⁵⁸), with a mean per-domain Spearman ρ of 0.1227 (median 0.1193) over 403,482 matched pairs (**Figure 3A**). The domains are not plant proteins, were never seen by the model, and no parameter was fitted to them, making this its broadest fully external test. Both quantities depend on residue identity by construction; that shared dependence is the mechanism under test, not a confound. The correlations are modest in size; what the test establishes is not accuracy of magnitude but direction, across an overwhelming majority of independent domains.

#### Whole proteomes withheld from training

The classifier was applied to three proteomes absent from the resource and from its training (peach, grapevine and poplar, none from a family in CropStabDB, though poplar shares an order with cassava), giving a pooled AUC of 0.9190 across 112,057 proteins (**Figure 3B**). Its target is a threshold on AlphaFold mean confidence, so this shows that the sequence signal for that threshold transfers to families the model has never seen.

#### Withholding relatives

Leave-one-species-out over 19 species gives AUC 0.9166 ± 0.0238 (**Figure 3C**, **Table S8**). Cross-validation with entire orthogroups withheld, grouped by connected components so no relative of a test protein remains in training, gives 0.8964 ± 0.0024 across five folds and 121,982 groups. Against a random split (0.9331 ± 0.0003) this locates the cost of generalisation: withholding relatives costs about 0.037 AUC, while restricting the data to representative isoforms changes almost nothing (0.8935 ± 0.0039), so the drop comes from removing orthologues rather than from isoform redundancy (**Figure 3D**).

### Rubisco activase: a case with measured thermostability

The enzyme whose engineered thermostability first demonstrated this lever provides an independent test of the score, and required no parameter choice on our part. Its three isoforms have had their thermostability measured *in vitro*: TaRca1-β is thermotolerant, with a thermal midpoint of 42 °C, while TaRca2-α and TaRca2-β are heat-labile, with thermal midpoints of 35 °C [7], and TaRca1-β accumulates preferentially under heat stress [25]. Applied blind to the wheat proteome, the score orders the isoforms the same way in all three subgenomes, giving Rca1 a mean CropStab score of 35.00 against 38.69 for Rca2 (lower is the stable direction). TaRca1-β was located not by name but through its published isoform-specific peptide KKELDEGKQTNADR, which matches a single sequence among the 107,137 wheat proteins (**Figure S5**).

The same protein allows the orthologue constraint to be tested against a measured outcome. Eleven substitutions, each the residue TaRca1-β carries at a column where the two isoforms differ, raise the thermal midpoint of the heat-labile isoform by more than 7 °C [7]. Of the 8,816 substitutions possible in the 464-residue wheat sequence stored here, 38 positions hold a residue that no stable member of the orthogroup shares, and ten of the eleven lie among them. Scoring those candidates, which the Atlas itself does not do here, places T405K, M407L and Q409E highest; these are the C-terminal set that alone accounts for 2.6 °C of the reported gain. Within the admitted candidates the score does not otherwise separate the published eleven (permutation P = 0.84), so the test supports the constraint rather than the ranking.

### Benchmark

So that other methods can be compared on identical data, CropStabDB ships a frozen two-task benchmark with public test sets, declared baseline directions and an evaluation script (**Figure 3E**). On the protein-level task the score reaches AUC 0.7237 against 0.7015 for the instability index, which also enters the score as SIS, and 0.6820 for mean predicted disorder. The substitution-level task uses the distributed ΔΔG values unchanged, scored as the signed mean within-domain Spearman correlation since a continuous target admits no ROC; each baseline’s expected sign was declared in advance, and the CropStab model reaches +0.4438, against three declared baselines (Δ TOP-IDP, Δ hydropathy [26] and Δ CropStab score, the plain difference in component score between the two residues) reported alongside in **Table S8**.

### Usage

A researcher arrives with one protein and leaves with a ranked shortlist of residues to change, at https://cropstab.org and without registration (**Figure 4A**–D). Proteins can be searched by identifier, description or gene symbol, browsed by species and class, and filtered on any component; each record shows its score, six component values, confidence layer and Atlas entries. The Atlas can also be browsed directly, and a protein compared against its orthologues across species.

**Figure 4.**
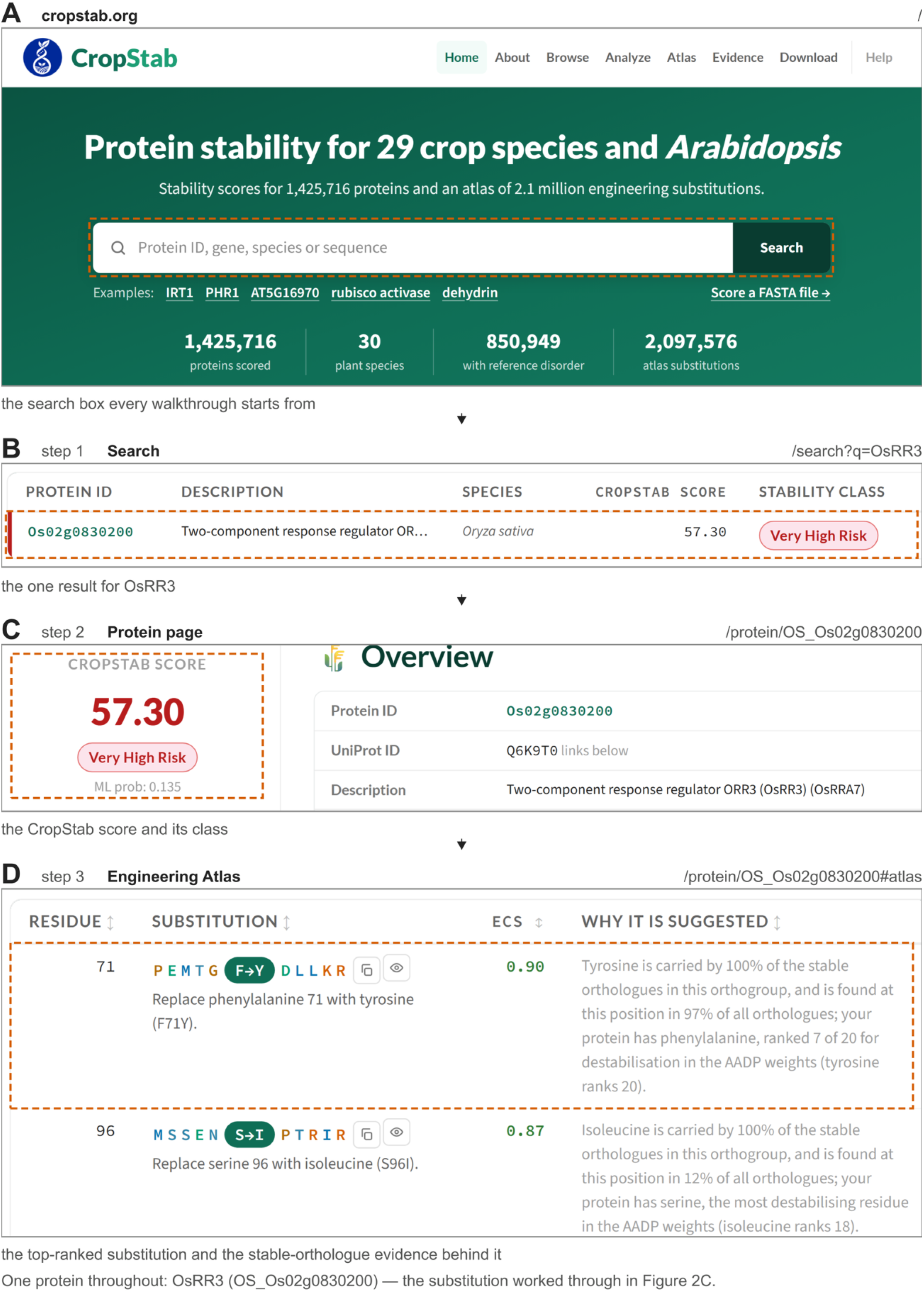
The three-step walkthrough at cropstab.org · **A** the home page and its search box · **B** step 1, the search result for OsRR3 · **C** step 2, the protein page with its score and class · **D** step 3, the Engineering Atlas entries for that protein

A sequence need not already be in the database. The analysis tool accepts a raw amino-acid sequence and returns its score and component profile, so the resource can be applied to a protein from any plant species, not only the 30 it contains; Atlas entries require an orthogroup, and so are available for proteins already in the database.

## Discussion

Protein stability is a sequence-encoded property that sets the conditions under which a protein folds, holds that fold and retains activity, and it therefore constrains photosynthesis, growth and yield in crops exposed to heat [3,5]. Engineering it has so far meant choosing a protein in advance and then building a library or an alignment for that protein alone [6,7]. CropStabDB inverts that order. The score ranks every protein in 30 plant proteomes and the Atlas proposes the residues, so choosing a target becomes a query rather than a commitment.

**Table S9** sets CropStabDB beside the resources closest to it. Measured data are more reliable but far narrower; predictors of melting temperature return one number and name no residue; structure-based force fields evaluate a substitution accurately but need a structure and need the substitution supplied; and experimental routes supply the orthologue constraint one protein at a time. Per-variant predictors answer a different question: FoldX [27] estimates the free-energy cost of a single substitution on a structure, and the ESM language model [28] scores variant likelihood from sequence, neither ranking whole proteomes. CropStabDB is the only resource in Table S9 that scores whole plant proteomes and proposes residues.

Ten features distinguish CropStabDB from these. (1) Coverage: all 1,425,716 proteins of 30 plant species are scored, not a sample. (2) Independence from structure: every value is computed from sequence, so coverage does not depend on a structural model existing. (3) Interpretability: a melting temperature is one number, whereas six components are six, and only one of their fifteen pairs is strongly correlated, so a score can be taken apart rather than only ranked. (4) Residue-level output: 2,097,576 candidate substitutions, each one a residue that a more stable orthologue already carries at the aligned column. (5) Locatability: every entry carries an 11-residue context string with the focal residue bracketed, and the 12,705 whose context is not unique are flagged. (6) Two confidence layers: AlphaFold per-residue confidence for 850,949 proteins, a sequence-based disorder layer for the remaining 574,767, and a record in every protein of which it has. (7) External validation: substitution-type scores match the direction of measured folding free energies in 842 of 926 domains, none of them plant proteins and none seen by the model, and in wheat Rubisco activase the orthologue constraint admits 38 of 464 positions, among them ten of the eleven substitutions measured to raise the thermal midpoint of the heat-labile isoform. (8) A frozen benchmark with public test sets, declared baseline directions and an evaluation script, so that any other method can be scored on the same data (**Table S8**). (9) Reproducibility: software versions are pinned, all parameters are deposited, and the serialised model reproduces its stored predictions across all 1,425,716 proteins. (10) Access without barrier: no registration, and a tool that scores a raw sequence from any plant species.

The score and the Atlas are not independent. An orthogroup’s stable and unstable members are set by fixed percentiles of the score distribution, so the Atlas exists because the score does, and the score continues where the Atlas stops: wheat Rubisco activase has no Atlas entries, so the isoform ranking above tests the score alone.

That the score tracks something biological can be seen outside the Atlas as well: it separates two classes of curated Arabidopsis nutrient genes in the direction their function suggests. Ion transporters score at a median of 29.58 against 54.20 for their transcription-factor regulators, a difference of 24.6 units across 27 curated genes (Mann–Whitney P = 2.2 × 10⁻⁵); the same regulators are unremarkable against 1,150 *Arabidopsis* transcription factors (P = 0.91, **Table S7**). The difference is recorded as an observation, and no mechanism is proposed for it.

The Atlas offers only what a stable relative already carries, which is why 172 of the 380 possible substitution types never appear, and experimental validation of individual proposals is the next step. CropStabDB will grow: further crop and wild-relative proteomes as the community asks for them, and an extended benchmark as measured stability data become available.

Protein stability determines whether a crop protein keeps working as temperature rises. CropStabDB scores every plant protein for it, and names the residues that a stable relative already uses.

## Data availability

CropStabDB is served from a dedicated virtual server maintained by the authors, requires no registration and sets no query limits; the corresponding author is responsible for it. The authors commit to maintaining the service for at least five years. Because a web service is a weaker guarantee than a file, the complete resource is openly deposited under a CC BY 4.0 licence at Zenodo (concept DOI https://doi.org/10.5281/zenodo.20551449, which always resolves to the latest version): the database, the released model, the input file names and checksums, the Engineering Atlas, the benchmark with its evaluation script, and the plotted values behind every figure. The resource therefore remains usable, and every figure reproducible, if the site ever becomes unavailable. Analysis code is available at https://github.com/Israfil-Hossen/CropStabDB under an MIT licence. Each release is identified by the checksum of its database file, recorded in the deposit alongside the scripts that produced it, and a future release will be deposited as a new version of the same record rather than replacing the current files. Corrections can be reported through the contact address on the site, and users are asked to cite both the article and the deposited version they used, since counts and scores change between releases.

## CRediT author statement

Md. Isrfil Hossen: Methodology, Software, Formal analysis, Data curation, Validation, Visualization, Writing – original draft. Yilin Pan: Formal analysis. Ping Lan: Conceptualization, Supervision, Funding acquisition, Project administration, Resources, Writing – review & editing. All authors have read and approved the final manuscript.

## Supplementary material

Supplementary Tables S1–S9 and Supplementary Figures S1–S5 accompany this article. The supplementary data archive is deposited with the resource at Zenodo.

## Acknowledgments

We express our gratitude to colleagues at the State Key Laboratory of Soil and Sustainable Agriculture for their comments on the manuscript.

