## Supplementary material for "CropStabDB: a protein stability resource and Engineering Atlas of orthologue-supported substitutions for 30 plant proteomes": CropStabDB_Tables_Supplementary.docx

**Table S1** Input files and md5 sums, per species

The input file for each species, with its md5 sum, the number of protein records it contains, the assembly recorded for it and the header evidence that assembly rests on. Provenance is settled by the deposited files and their checksums rather than by a release string. No file size is given: the input FASTAs are not redistributed with the analysis code, so a size could only be copied from a note rather than measured.

| **Species** | **Input file** | **md5 of the input file** | **Protein records in the file** | **md5 provenance** | **Assembly (as recorded)** | **Header evidence recorded for that assembly** | **Download source** |
| --- | --- | --- | --- | --- | --- | --- | --- |
| *A. thaliana* | arabidopsis.fa | ba30f37dd34024c31ce612608f7e6fce | 27,306 | recorded twice, agreeing; recomputed and matched 2026-09-28 | TAIR10 | AGI identifiers AT#G#####.# (+ ATCG/ATMG organelle genes) | not recorded |
| *B. napus* | brassica_napus.fa | 59cb3b913fc36bff32948aef3628da75 | 101,040 | recorded twice, agreeing; recomputed and matched 2026-09-28 | AST_PRJEB5043_v1 | assembly named in every Ensembl Plants pep header | Ensembl Plants |
| *B. oleracea* | brassica_oleracea.fa | b76ffc5c480f58f6af99cdd74a42b64e | 59,220 | recorded twice, agreeing; recomputed and matched 2026-09-28 | BOL | assembly named in every Ensembl Plants pep header | Ensembl Plants |
| *B. rapa* | brassica_rapa.fa | f5eb7539522c43d75c1412e55458be58 | 41,025 | recorded twice, agreeing; recomputed and matched 2026-09-28 | Brapa_1.0 | assembly named in every Ensembl Plants pep header | Ensembl Plants |
| *B. vulgaris* | beta_vulgaris.fa | 69e1750ba1224d36eab018cd060e6d29 | 28,721 | recorded twice, agreeing; recomputed and matched 2026-09-28 | RefBeet-1.2.2 | assembly named in every Ensembl Plants pep header | Ensembl Plants |
| *C. annuum* | capsicum_annuum.fa | d65a5e40efbd5bbf5d60c412d55f2a42 | 35,845 | recorded twice, agreeing; recomputed and matched 2026-09-28 | ASM51225v2 | assembly named in every Ensembl Plants pep header | Ensembl Plants |
| *C. canephora* | coffea_canephora.fa | f9db9859af77a55a5b478efabe6a785d | 25,574 | recorded twice, agreeing; recomputed and matched 2026-09-28 | AUK_PRJEB4211_v1 | assembly named in every Ensembl Plants pep header | Ensembl Plants |
| *C. sativus* | cucumis_sativus.fa | 1af6e975ea9e40896091aa93271f6b24 | 23,780 | recorded twice, agreeing; recomputed and matched 2026-09-28 | ASM407v2 | assembly named in every Ensembl Plants pep header | Ensembl Plants |
| *D. carota* | daucus_carota.fa | 4a54b5347cf7d1508612e2df834dca0f | 32,109 | recorded twice, agreeing; recomputed and matched 2026-09-28 | ASM162521v1 | assembly named in every Ensembl Plants pep header | Ensembl Plants |
| *E. coracana* | eleusine_coracana.fa | a3aee7922d7a3a38e173d1548bedac59 | 57,180 | recorded twice, agreeing; recomputed and matched 2026-09-28 | Eleusine coracana PR202 (NCBI; version not stated in the file) | NCBI protein accessions GJM/GJN, locus tags PR202_ | NCBI |
| *G. max* | soybean.fa | 3829f15adef9d1e8d0a15fc4e6549b16 | 55,653 | recorded twice, agreeing; recomputed and matched 2026-09-28 | Glycine max Wm82.a2 (v2.0 or v2.1 - not decidable from identifiers) | GLYMA_ gene identifiers; INSDC protein accessions KRH/KRG/RCW | not recorded |
| *G. raimondii* | gossypium_raimondii.fa | 9b5b50c23b997640ea951ec48429bf58 | 78,371 | recorded twice, agreeing; recomputed and matched 2026-09-28 | Graimondii2_0_v6 | assembly named in every Ensembl Plants pep header | Ensembl Plants |
| *H. annuus* | helianthus_annuus.fa | 6c814533c717205a5c9253bb6f354da7 | 70,864 | recorded twice, agreeing; recomputed and matched 2026-09-28 | HanXRQr2.0-SUNRISE | assembly named in every Ensembl Plants pep header | Ensembl Plants |
| *H. vulgare* | barley.fa | 0cd7dbd39b5fdd69135c93a0fd003327 | 35,825 | recorded twice, agreeing; recomputed and matched 2026-09-28 | MorexV3 | HORVU.MOREX.r3 identifiers | not recorded |
| *M. acuminata* | musa_acuminata.fa | 91471b199d85332a2248f6290659a086 | 45,589 | recorded twice, agreeing; recomputed and matched 2026-09-28 | Musa_acuminata_v2 | assembly named in every Ensembl Plants pep header | Ensembl Plants |
| *M. esculenta* | manihot_esculenta.fa | a0cacbec94127b43bcbf541159b018bc | 59,149 | recorded twice, agreeing; recomputed and matched 2026-09-28 | M.esculenta_v8 | assembly named in every Ensembl Plants pep header | Ensembl Plants |
| *M. truncatula* | medicago_truncatula.fa | e261b3152be8ab210dc270d7f6e10ad2 | 57,585 | recorded twice, agreeing; recomputed and matched 2026-09-28 | MedtrA17_4.0 | assembly named in every Ensembl Plants pep header | Ensembl Plants |
| *O. sativa* | rice.fa | a6fdde8344c0f53c29b5296508521d8d | 35,359 | recorded twice, agreeing; recomputed and matched 2026-09-28 | IRGSP-1.0 | RAP-DB identifiers Os##t#######-## (35,265) + 94 GenBank organelle CDS | not recorded |
| *P. hallii* | panicum_hallii_fil2.fa | e2d979899f2f227027662726a62c9a4e | 44,192 | recorded twice, agreeing; recomputed and matched 2026-09-28 | PHallii_v3.1 | assembly named in every Ensembl Plants pep header | Ensembl Plants |
| *P. sativum* | pisum_sativum.fa | d8d7f7ec5de34e7f9f1c36a6c266a791 | 56,770 | recorded twice, agreeing; recomputed and matched 2026-09-28 | Pisum_sativum_v1a | assembly named in every Ensembl Plants pep header | Ensembl Plants |
| *P. vulgaris* | phaseolus_vulgaris.fa | d77303a92c137aacd4c30d625ff9a472 | 32,720 | recorded twice, agreeing; recomputed and matched 2026-09-28 | PhaVulg1_0 | assembly named in every Ensembl Plants pep header | Ensembl Plants |
| *S. bicolor* | sorghum.fa | 7c3fec7c269f3d7235dc7cfab7eec071 | 33,878 | recorded twice, agreeing; recomputed and matched 2026-09-28 | Sorghum bicolor v3 (NCBIv3) | SORBI_3 gene identifiers; INSDC protein accessions OQU/KXG/EES | not recorded |
| *S. cereale* | secale_cereale.fa | 59ce689db9ed9c5403a577c32562a65c | 34,441 | recorded twice, agreeing; recomputed and matched 2026-09-28 | Rye_Lo7_2018_v1p1p1 | assembly named in every Ensembl Plants pep header | Ensembl Plants |
| *S. italica* | setaria_italica.fa | 3b25580f05bef5fa70934888c37536e2 | 41,023 | recorded twice, agreeing; recomputed and matched 2026-09-28 | Setaria_italica_v2.0 | assembly named in every Ensembl Plants pep header | Ensembl Plants |
| *S. lycopersicum* | solanum_lycopersicum.fa | fcae567e6cbf98adc8869a60f376982b | 34,429 | recorded twice, agreeing; recomputed and matched 2026-09-28 | SL3.0 | assembly named in every Ensembl Plants pep header | Ensembl Plants |
| *S. tuberosum* | solanum_tuberosum.fa | 528bf162ff45c9c95f18b42051a65a1c | 56,210 | recorded twice, agreeing; recomputed and matched 2026-09-28 | SolTub_3.0 | assembly named in every Ensembl Plants pep header | Ensembl Plants |
| *T. aestivum* | wheat.fa | b6b40088e2e98e34dd299817699396df | 107,137 | recorded twice, agreeing; recomputed and matched 2026-09-28 | IWGSC RefSeq v1.0 assembly (annotation v1.0 vs v1.1 not decidable from identifiers) | TraesCS#?##G###### identifiers | not recorded |
| *V. radiata* | vigna_radiata.fa | b3853d06815fbc8490639dadbfa72544 | 23,181 | recorded twice, agreeing; recomputed and matched 2026-09-28 | Vradiata_ver6 | assembly named in every Ensembl Plants pep header | Ensembl Plants |
| *V. unguiculata* | vigna_unguiculata.fa | 9c7f4b150378dab9bdbcad7b34216d9e | 54,346 | recorded twice, agreeing; recomputed and matched 2026-09-28 | ASM411807v1 | assembly named in every Ensembl Plants pep header | Ensembl Plants |
| *Z. mays* | maize.fa | 005a2afdcb8b80c886775a8c0b276e8a | 39,756 | recorded twice, agreeing; recomputed and matched 2026-09-28 | Zm-B73-REFERENCE-NAM-5.0 | Zm00001eb identifiers | not recorded |
| Total (30 species) |  |  | 1,428,278 |  |  |  |  |

^a^ Source: species_provenance.tsv (md5 d5b8b4543235f839f9b2848d2f5e5f6a).

^b^ The header evidence column reports what was observed in each file's headers when the provenance record was made; it is not re-read in this run.

^c^ 7 species have no recorded download source and say so.

^d^ Every md5 is recorded in two independent places that agree for all 30 species, and all 30 were recomputed and matched on 2026-09-28 by VERIFICATION_SCRIPTS/tables/TS1_input_files.py, which reads the retained local copy of each proteome and takes the md5 of the decompressed stream. The retained copies are not part of the deposit: the providers' data-use terms differ and no right to republish them is claimed, so the record carries each file's name and checksum and not the file. The md5 provenance column states this per row rather than leaving it to be assumed.

*Built by VERIFICATION_SCRIPTS/tables/TS4_input_files.py.*

**Table S2** Candidate features and selection

The deposited candidate feature set and the selection that produced the released model's 64 features. The deposited lists show three features removed at the first step and seven at the second. The starting set is not deposited as a named list; the earliest deposited list holds 74 features, so the selection from 74 to 64 is what this table lets a reader check.

| **Feature** | **Category (from the feature dictionary)** | **Selected** | **Stage** | **Note** |
| --- | --- | --- | --- | --- |
| aa_A | Amino-acid composition | yes | retained | kept through both selection stages |
| aa_C | Amino-acid composition | yes | retained | kept through both selection stages |
| aa_D | Amino-acid composition | yes | retained | kept through both selection stages |
| aa_E | Amino-acid composition | yes | retained | kept through both selection stages |
| aa_F | Amino-acid composition | yes | retained | kept through both selection stages |
| aa_G | Amino-acid composition | yes | retained | kept through both selection stages |
| aa_H | Amino-acid composition | yes | retained | kept through both selection stages |
| aa_I | Amino-acid composition | yes | retained | kept through both selection stages |
| aa_K | Amino-acid composition | yes | retained | kept through both selection stages |
| aa_L | Amino-acid composition | yes | retained | kept through both selection stages |
| aa_M | Amino-acid composition | yes | retained | kept through both selection stages |
| aa_N | Amino-acid composition | yes | retained | kept through both selection stages |
| aa_P | Amino-acid composition | yes | retained | kept through both selection stages |
| aa_Q | Amino-acid composition | yes | retained | kept through both selection stages |
| aa_R | Amino-acid composition | yes | retained | kept through both selection stages |
| aa_S | Amino-acid composition | yes | retained | kept through both selection stages |
| aa_T | Amino-acid composition | yes | retained | kept through both selection stages |
| aa_V | Amino-acid composition | yes | retained | kept through both selection stages |
| aa_W | Amino-acid composition | yes | retained | kept through both selection stages |
| aa_Y | Amino-acid composition | yes | retained | kept through both selection stages |
| abox_count | Repeats and low complexity | yes | retained | kept through both selection stages |
| aliphatic_index | Global physicochemical | yes | retained | kept through both selection stages |
| aromatic_percent | Amino-acid composition | yes | retained | kept through both selection stages |
| c_terminal_aa_enc | Terminal residues | yes | retained | kept through both selection stages |
| charge_at_pH7 | Global physicochemical | yes | retained | kept through both selection stages |
| charged_percent | Amino-acid composition | yes | retained | kept through both selection stages |
| coil_fraction | Secondary-structure propensity | yes | retained | kept through both selection stages |
| dbox_count | Degradation motifs | yes | retained | kept through both selection stages |
| extinction_coefficient | Global physicochemical | yes | retained | kept through both selection stages |
| gravy | Global physicochemical | yes | retained | kept through both selection stages |
| has_chloroplast_tp | Localization and targeting signals | yes | retained | kept through both selection stages |
| has_er_retention | Localization and targeting signals | yes | retained | kept through both selection stages |
| has_gpi_anchor | Localization and targeting signals | yes | retained | kept through both selection stages |
| has_mitochondrial_tp | Localization and targeting signals | yes | retained | kept through both selection stages |
| has_nls | Localization and targeting signals | yes | retained | kept through both selection stages |
| has_pest | Degradation motifs | yes | retained | kept through both selection stages |
| has_signal_peptide | Localization and targeting signals | yes | retained | kept through both selection stages |
| helix_fraction | Secondary-structure propensity | yes | retained | kept through both selection stages |
| hydrophobic_percent | Amino-acid composition | yes | retained | kept through both selection stages |
| isoelectric_point | Global physicochemical | yes | retained | kept through both selection stages |
| kenbox_count | Degradation motifs | yes | retained | kept through both selection stages |
| lcr_fraction | Repeats and low complexity | yes | retained | kept through both selection stages |
| length | Global physicochemical | yes | retained | kept through both selection stages |
| max_repeat_length | Repeats and low complexity | yes | retained | kept through both selection stages |
| mean_disorder_score | Intrinsic disorder | yes | retained | kept through both selection stages |
| molecular_weight_kDa | Global physicochemical | yes | retained | kept through both selection stages |
| n_terminal_aa_enc | Terminal residues | yes | retained | kept through both selection stages |
| nglyc_sites | Post-translational modification | yes | retained | kept through both selection stages |
| pest_count | Degradation motifs | yes | retained | kept through both selection stages |
| pest_density | Degradation motifs | yes | retained | kept through both selection stages |
| pest_max_length | Degradation motifs | yes | retained | kept through both selection stages |
| pest_total_length | Degradation motifs | yes | retained | kept through both selection stages |
| polar_percent | Amino-acid composition | yes | retained | kept through both selection stages |
| predicted_localization_enc | Global physicochemical | yes | retained | kept through both selection stages |
| ptm_density | Post-translational modification | yes | retained | kept through both selection stages |
| repeat_fraction | Repeats and low complexity | yes | retained | kept through both selection stages |
| sheet_fraction | Secondary-structure propensity | yes | retained | kept through both selection stages |
| tm_domain_count | Localization and targeting signals | yes | retained | kept through both selection stages |
| total_destruction_signals | Degradation motifs | yes | retained | kept through both selection stages |
| total_phospho_sites | Post-translational modification | yes | retained | kept through both selection stages |
| total_repeat_residues | Repeats and low complexity | yes | retained | kept through both selection stages |
| total_repeats_3plus | Repeats and low complexity | yes | retained | kept through both selection stages |
| total_repeats_5plus | Repeats and low complexity | yes | retained | kept through both selection stages |
| ubiq_sites | Post-translational modification | yes | retained | kept through both selection stages |
| aromaticity | dropped before release; no row in the deposited dictionary | no | removed at stage 2 (71 to 64) | dropped in the second pass |
| charge_class_enc | dropped before release; no row in the deposited dictionary | no | removed at stage 2 (71 to 64) | dropped in the second pass |
| cysteine_percent | dropped before release; no row in the deposited dictionary | no | removed at stage 1 (74 to 71) | dropped in the first pass |
| glycine_percent | dropped before release; no row in the deposited dictionary | no | removed at stage 1 (74 to 71) | dropped in the first pass |
| hydrophobicity_class_enc | dropped before release; no row in the deposited dictionary | no | removed at stage 2 (71 to 64) | dropped in the second pass |
| is_high_ptm | dropped before release; no row in the deposited dictionary | no | removed at stage 2 (71 to 64) | dropped in the second pass |
| kferq_count | dropped before release; no row in the deposited dictionary | no | removed at stage 2 (71 to 64) | dropped in the second pass |
| kferq_density | dropped before release; no row in the deposited dictionary | no | removed at stage 2 (71 to 64) | dropped in the second pass |
| proline_percent | dropped before release; no row in the deposited dictionary | no | removed at stage 1 (74 to 71) | dropped in the first pass |
| size_class_enc | dropped before release; no row in the deposited dictionary | no | removed at stage 2 (71 to 64) | dropped in the second pass |

^a^ Lists: feature_names_FINAL_74.txt (md5 ab39d358fd259dd89dc6de4951883208), _71.txt, _64.txt (md5 a7cb47cd104814c736891c5283a08a2c).

^b^ Category is read from the feature dictionary and is a cross-check only; the lists themselves come from the deposited feature files.

^c^ No AUC is given for any feature set: the 64-feature AUC is not recorded anywhere in the deposit and is not reconstructed here.

*Built by VERIFICATION_SCRIPTS/tables/TS1_features.py.*

**Table S3** The NDS lookup table

The N-terminal Descriptor Score lookup, published in full as a parameter of the CropStab score. The residue is the N-terminal one after initiator-methionine excision. The values are quantised to 0.1 steps and represent a hand-assigned ordering, not a measured scale; no citation is given because none exists. The lookup is published so that it can be inspected and replaced. NDS carries weight 0.03, the smallest of the six components.

| **N-terminal residue (one letter)** | **NDS value** |
| --- | --- |
| G | 0.0 |
| M | 0.0 |
| P | 0.0 |
| V | 0.0 |
| A | 0.1 |
| S | 0.1 |
| I | 0.2 |
| T | 0.2 |
| C | 0.3 |
| L | 0.4 |
| H | 0.5 |
| F | 0.6 |
| W | 0.6 |
| Y | 0.6 |
| D | 0.7 |
| E | 0.7 |
| K | 0.7 |
| N | 0.7 |
| Q | 0.7 |
| R | 0.8 |
| any residue not listed | 0.3 (default) |

^a^ Read from the released scorer cropstab_score.py (md5 fb29f5348ae918cade833787a28c182e).

^b^ Weight from weights_v2.json (md5 2f2593f90d79311cfae8011d0a570324).

^c^ Table S5 reports the NDS component at AUC 0.4766, below chance.

*Built by VERIFICATION_SCRIPTS/tables/TS5_nds_lookup.py.*

**Table S4** Pairwise correlation between the six components

Spearman correlation between the six component scores over all 1,425,716 scored proteins, all 15 unique pairs. Exactly one pair exceeds |rho| = 0.5. The components are therefore not one quantity measured six times.

| **Component A** | **Component B** | **Spearman rho** | **Above \|rho\| 0.5** |
| --- | --- | --- | --- |
| AADP | SIS | +0.5531 | yes |
| SIS | SDS | +0.4907 | no |
| AADP | PTMS | +0.3562 | no |
| SDS | LAI | +0.3324 | no |
| AADP | SDS | +0.2926 | no |
| SIS | LAI | +0.2893 | no |
| AADP | LAI | +0.2678 | no |
| SDS | PTMS | +0.1710 | no |
| LAI | PTMS | +0.1710 | no |
| LAI | NDS | +0.0319 | no |
| SIS | PTMS | +0.0305 | no |
| AADP | NDS | -0.0271 | no |
| SDS | NDS | +0.0212 | no |
| PTMS | NDS | -0.0134 | no |
| SIS | NDS | -0.0012 | no |

^a^ Recomputed in this run from cropstabdb_v21i.sqlite, n = 1,425,716; the rank transform uses every protein, with no sampling.

^b^ NDS and LAI are discrete lookups, taking 20 and a handful of values respectively, so their rho carries more ties than the continuous components. That is a property of how those two components are defined.

*Built by VERIFICATION_SCRIPTS/tables/TS6_component_correlation.py.*

**Table S5** Component AUC, skill and weight

The six component scores, with the area under the ROC curve against the AlphaFold-derived target the classifier is trained on, the skill each contributes, and the weight it receives. AUC is reported in the declared direction with no direction correction, so NDS appears at 0.4766, below chance. Weight is proportional to |2 x AUC - 1|; because that expression takes an absolute value, a component below chance still receives a positive weight, and NDS's 0.03 is therefore not evidence that it discriminates. The final row gives the four class cut points and the proteome median.

| **Component** | **What it measures** | **AUC vs the AlphaFold-derived target, declared direction** | **Skill = \|2 x AUC - 1\|** | **Weight** | **Weight (%)** |
| --- | --- | --- | --- | --- | --- |
| AADP | Amino-acid composition weighted by the pLDDT-model SHAP direction | 0.7154 | 0.4307 | 0.27 | 27% |
| SIS | ProtParam instability index, logistic-scaled | 0.7044 | 0.4088 | 0.25 | 25% |
| SDS | Mean TOP-IDP disorder propensity over a 21-residue window, rescaled to 0-1 | 0.6925 | 0.3850 | 0.24 | 24% |
| LAI | Compartment weight for the predicted subcellular localisation | 0.5870 | 0.1739 | 0.11 | 11% |
| PTMS | Phosphorylation, N-glycosylation and ubiquitination sites per 100 aa | 0.5823 | 0.1645 | 0.10 | 10% |
| NDS | Class of the N-terminal residue after initiator-Met excision, from a fixed lookup | 0.4766 (below chance) | 0.0468 | 0.03 | 3% |
| Sum |  |  | 1.6098 | 1.00 | 100% |
| Class cut points | Classes are quintiles of the scored proteome (1,425,716 proteins) | 36.99 / 41.82 / 46.62 / 52.81 | proteome median 44.14 |  |  |

^a^ n = 851,077 proteins with a matched AlphaFold model; target is mean pLDDT < 70.

^b^ Recomputed here from master_v2.csv (md5 fa7e0f63e7aeff7b75590c87fbb1c41d) and alphafold_matched_v2.csv (md5 9f25c2f996a361f17c1b60e1bd15cfac).

^c^ Machine-readable records elsewhere in the deposit store NDS as 0.5234, the direction-corrected form of the same measurement (0.5234 = 1 - 0.4766). No weight is affected: the skill is 0.0468 either way.

^d^ The NDS lookup table is published in full as Table S3. It is a parameter of the score, not a measured scale, and carries no citation because none exists.

*Built by VERIFICATION_SCRIPTS/tables/TS2_components.py.*

**Table S6** Atlas column dictionary

Every column of the released Engineering Atlas, with its meaning, the number of distinct values, the range, and the null count, all read from the deposited file. No column contains a null. Four columns carry names that mislead and are corrected in bold: `position` is an alignment column and not a residue number, `aadp_delta` (database `delta_shap`) is not a SHAP value, `current_is_minority` (database `current_fraction`) is a three-valued flag and not a fraction, and `component_fix` is a cut-point rather than a component identity.

| **Column** | **Meaning** | **Distinct values** | **Range or values** | **Nulls** |
| --- | --- | --- | --- | --- |
| protein_id | Protein identifier, as in the proteins table | 51,440 | 8-14 chars | 0 |
| species | Species key used throughout the deposit | 30 | 11-5 chars | 0 |
| orthogroup | OrthoFinder orthogroup | 6,223 | 9-9 chars | 0 |
| position | **NOT a residue number. This is the ALIGNMENT COLUMN index in the orthogroup MSA (1-based). residue_position is the number to use. Confusing the two produced the discarded coverage figure that was withdrawn.** Database name: alignment_column | 2,799 | 1 to 5550 | 0 |
| from_aa | Current residue, one letter | 19 | 1-1 chars | 0 |
| to_aa | Proposed residue, one letter | 19 | 1-1 chars | 0 |
| CropStab_Score_v2 | The protein's CropStab score, repeated on every row | 13,010 | 52.806 to 81.22 | 0 |
| ortholog_score | Share of the orthogroup's stable members carrying the proposed residue at this column. Database name: n_stable | 909 | 0.012 to 1 | 0 |
| aadp_delta | **NOT a SHAP value. It is the AADP weight difference: AA_W[from_aa] - AA_W[to_aa].** Database name: delta_shap | 195 | -0.00095 to 0.041 | 0 |
| aadp_agrees | Cut-point on aadp_delta: 1.0 when > 0.001, else 0.5. 0.0 never occurs | 2 | 0.5 to 1 | 0 |
| conservation_of_to | Frequency of the proposed residue within its alignment column. A LOCAL column frequency, not an evolutionary conservation analysis | 971 | 0.007 to 0.993 | 0 |
| current_is_minority | **NOT a fraction. A three-valued flag: 1.0 when the current residue's column frequency is below 0.30, 0.5 when below 0.50, else 0.0.** Database name: current_fraction, which is what makes it misleading | 3 | 0 to 1 | 0 |
| pest_disruption | Flag: 1.0 PEST to hydrophobic, 0.5 PEST to non-PEST, else 0.0 | 3 | 0 to 1 | 0 |
| component_fix | **A second, coarser cut-point on aadp_delta: 1.0 if aadp_delta > 0.005 else 0.5. It is NOT a component identity. No Atlas column records which component a substitution acts on.** | 2 | 0.5 to 1 | 0 |
| ECS | Engineering Confidence Score: 0.30 x ortholog_score + 0.20 x aadp_agrees + 0.15 x conservation_of_to + 0.15 x current_is_minority + 0.10 x pest_disruption + 0.10 x component_fix | 5,984 | 0.4 to 0.999 | 0 |
| residue_position | Residue number in the processed sequence, 1-based. This is the number to use | 2,265 | 1 to 3778 | 0 |
| position_not_unique_v21 | True when the residue number is not uniquely determined by remapping | 2 | 0 to 1 | 0 |
| context | Five residues each side, target bracketed, ^/$ for a terminus | 289,199+ | 13-9 chars | 0 |
| protein_length | Length of the processed sequence | 1,950 | 30 to 3789 | 0 |
| remap_status_v21 | Status of the alignment-to-FASTA remapping | 1 | ok | 0 |
| db_length | Length stored in the proteins table | 1,950 | 30 to 3789 | 0 |
| position_unique | 1 when the residue number is uniquely determined | 2 | 0 to 1 | 0 |
| position_not_unique | Complement of position_unique | 2 | 0 to 1 | 0 |

^a^ Source: engineering_atlas_v21_crosswalk_full.csv (md5 609062af95c57123ad187ab1a0262812), 2,097,576 rows.

^b^ No Atlas column records which of the six components a substitution acts on.

^c^ db_length equals protein_length on every retained row, and remap_status_v21 is ok throughout.

*Built by VERIFICATION_SCRIPTS/tables/TS3_atlas_columns.py.*

**Table S7** Quality-control and reference statistics

Quality-control and reference statistics. These values are quoted in the text and are collected here so each can be checked in one place. Every row was recomputed from the release database or the deposited files in the run that produced this table, except the two marked "not recomputed here", which say why. The rows are measurements of how the resource behaves, not results: no figure is drawn from them, and the pLDDT row is a reference rather than a benchmark for the reason its scope column gives.

| **Quantity** | **Value** | **n** | **Recomputed in this run by** | **Scope and provenance** |
| --- | --- | --- | --- | --- |
| Proteins carrying a sequence-based disorder layer | 574,767 | 1,425,716 proteins | TS7_quality_control.py | 1,425,716 total minus 850,949 with an AlphaFold layer = 574,767. Counted as disorder_source = 'predicted_topidp' in the release database, which equals that subtraction exactly. |
| Largest single cell of the substitution matrix (serine to alanine) | 54,984 | 2,097,576 Atlas entries | TS7_quality_control.py | Counted directly; the matrix is drawn in Supplementary Figure S4. Of the 380 possible ordered substitution types, 208 are ever proposed. |
| Atlas entries whose 11-residue context is not unique within its protein | 12,705 | 2,097,576 Atlas entries | TS7_quality_control.py | Counted from the position_not_unique flag stored on every Atlas row. |
| Proteins in the orthogroup set used for the behaviour checks | 1,156,886 | 60,214 orthogroups, 30 species | TS7_quality_control.py | Representative isoforms only, 'unassigned' excluded, orthogroups with at least two members. Quality control, not a result; no figure is drawn from it. |
| Agreement between orthologue scores across species (Spearman rho) | +0.7244 | 45,043 one-per-orthogroup pairs | TS7_quality_control.py | One pair drawn per orthogroup, two species picked at random, RandomState(0). Quality control, not a result; no figure is drawn from it. |
| The same measurement on shuffled pairs (control) | +0.0077 | 45,043 pairs | TS7_quality_control.py | The same scores with one side permuted, same seed. A control for the row above. |
| Reference comparison: CropStab score against AlphaFold mean pLDDT < 70 (ROC AUC) | 0.7316 | 850,949 proteins, 363,734 below the threshold | TS7_quality_control.py | A REFERENCE, NOT A BENCHMARK. One component, AADP, is weighted using pLDDT, so this comparison is partly circular. That is why it is not a row of Table 2. |
| Groups withheld whole in the orthogroup-blocked cross-validation | 121,982 | 5 folds, 851,077 labelled proteins | not recomputed here | The grouping method reproduces - connected components joining proteins that share an orthogroup or a gene, singletons included - but the count does not. Recomputing it over the 850,949 labelled proteins in the release database gives 122,304. The recorded figure was measured on the 851,077-protein model set: 128 proteins differ, and that moves the component count by 322. The recorded value is quoted, not the recomputed one, and the discrepancy is stated rather than absorbed. |
| Effect of setting the NDS component weight to zero: change in ROC AUC | +0.0004 | 850,949 proteins with an AlphaFold layer | TS7_quality_control.py | The remaining five weights are renormalised to the same total. Against the same pLDDT < 70 reference target as the row above, so the same circularity applies. |
| Effect of setting the NDS component weight to zero: rank correlation with the released score | 0.9996 | 1,425,716 proteins | TS7_quality_control.py | Spearman correlation between the released score and the same score with NDS removed. |
| Proteins from which one or more characters were removed during cleaning | 3,687 | 1,425,716 retained of 1,428,278 input records | TS7_quality_control.py | The cleaning rule of 03_provenance/regenerate_processed.py replayed over all 30 input proteomes: upper-case, drop '*' and 'X', keep only the 20 standard residues, keep the protein at 30 residues or more. The proteomes are RETAINED LOCALLY but NOT DEPOSITED - the providers' terms differ and no right to republish them is claimed - so a reader repeats this by downloading each file from its provider, which species_provenance.tsv identifies by name and md5. |
| Curated Arabidopsis nutrient transporters: median CropStab score | 29.58 | 17 curated genes | TS7_quality_control.py | Curated by gene identifier in the release database's nutrient_genes table, role = 'transporter'; scores read from the proteins table, not from the rounded values stored beside the gene. Quality control, not a result. |
| Curated Arabidopsis nutrient transcription-factor regulators: median CropStab score | 54.20 | 10 curated genes | TS7_quality_control.py | role = 'regulator' in the same table. The difference between the two medians is 24.6145 score units, reported in the text as 24.6; two-sided Mann-Whitney P = 2.2e-05 over the 27 genes carrying either role. Three further curated genes carry role 'other' and enter no statistic. |
| NEGATIVE CONTROL - the same regulators against every Arabidopsis transcription factor (two-sided Mann-Whitney P) | 0.91 | 10 regulators against 1,150 transcription factors | TS7_quality_control.py | The regulators score low for Arabidopsis proteins generally, but they are UNREMARKABLE among transcription factors: this test does not reject the null. It bounds the separation reported in the Discussion - the score is not shown to single these regulators out from transcription factors as a class - and it is reported whether or not it flatters the result. Transcription factors are those of the tf_advisor table. |
| Orthogroups in the release database | 63,185 | 1,316,830 proteins carrying an orthogroup | TS7_quality_control.py | Distinct values of proteins.orthogroup excluding the literal 'unassigned', which 108,886 proteins carry. Counting the distinct values without that exclusion gives 63,186, one more, and that is the only difference between the two figures. Of these, 6,223 hold both a stable and an unstable member and so contribute to the Engineering Atlas. |

^a^ Release database CroStab_Final/v2_release/07_release/cropstabdb_v21i.sqlite, md5 verified before reading.

^b^ Component weights read from 04_model/02_components/weights_v2.json (md5 2f2593f90d79311cfae8011d0a570324).

^c^ ROC AUC is computed from ranks, with no model fitted and no threshold chosen.

*Built by VERIFICATION_SCRIPTS/tables/TS7_quality_control.py.*

**Table S8** Validation and benchmark results

Validation and benchmark results. Validation is listed in decreasing order of independence: the comparison against measured folding free energies depends on neither a structural model nor the resource's own residue numbering, and is reported first. Benchmark rows give each method's result on the two frozen tasks, with the direction each was expected to score in declared before scoring.

| **Part** | **What is tested** | **Method** | **Metric** | **Result** | **n** | **Direction declared in advance** | **Source** |
| --- | --- | --- | --- | --- | --- | --- | --- |
| A. Validation | Agreement with measured folding free energies (non-crop domains) | Engineering Atlas substitution-type mean ECS; no model fitted | Per-domain Spearman rho vs ddG; share of domains positive | 842 of 926 domains positive (90.9%); mean rho 0.1227, median 0.1193; P = 1.91e-158 | 926 domains, 403,482 matched pairs | positive rho expected | Tsuboyama et al. 2023; VERIFICATION_SCRIPTS/78 (crosswalk) |
| A. Validation | Three proteomes absent from CropStabDB (grapevine, poplar, peach) | Released random forest, 64 features (6_MODEL.joblib) | Pooled ROC AUC against AlphaFold mean pLDDT < 70 | 0.9190 (peach 0.9264, grapevine 0.9153, poplar 0.9148) | 112,057 proteins, 3 species | higher = more disordered | 06_validation/holdout_summary.csv; 05_benchmark/evaluate_v2.py |
| A. Validation | Held-out species | Released model configuration, all isoforms | Leave-one-species-out ROC AUC, mean +- SD | 0.9166 +- 0.0238 (range 0.8428-0.9551) | 19 species held out in turn | higher = more disordered | 06_validation/loso_all_isoforms.csv |
| A. Validation | Held-out orthogroups | Released model configuration, all isoforms | 5-fold cross-validation ROC AUC, whole orthogroups withheld, mean +- SD | 0.8964 +- 0.0024 | 5 folds, 851,077 test proteins, 121,982 groups | higher = more disordered | 06_validation/cv_all_isoforms_orthogroup_withheld.csv |
| B. Benchmark | Task 1 and Task 2 | CropStab model (released classifier / 64-feature deltas) | Task 1 pooled ROC AUC \| Task 2 signed mean within-domain Spearman rho | Task 1 0.9190 \| Task 2 +0.4438 | Task 1 112,057 proteins \| Task 2 150 domains, 164,070 mutations | +1 \| +1 | 05_benchmark/evaluate_v2.py on the deposited baseline files |
| B. Benchmark | Task 1 and Task 2 | CropStab score / delta CropStab score | Task 1 pooled ROC AUC \| Task 2 signed mean within-domain Spearman rho | Task 1 0.7237 \| Task 2 -0.2706 | Task 1 112,057 proteins \| Task 2 150 domains | +1 \| -1 | 05_benchmark/benchmark_spec_v2.json |
| B. Benchmark | Task 1 and Task 2 | Mean predicted disorder (TOP-IDP, 21-residue window) / delta TOP-IDP | Task 1 pooled ROC AUC \| Task 2 signed mean within-domain Spearman rho | Task 1 0.6820 \| Task 2 -0.3849 | Task 1 112,057 proteins \| Task 2 150 domains | +1 \| -1 | spec; Task 2 recomputed from the deposited TOP-IDP scale |
| B. Benchmark | Task 2 only | Delta hydropathy (GRAVY, Kyte-Doolittle) | Signed mean within-domain Spearman rho | Task 2 +0.3343 | 150 domains, 164,070 mutations | +1 | spec; recomputed from the deposited Kyte-Doolittle scale |
| B. Benchmark | Task 1 only | Instability index (ProtParam) | Pooled ROC AUC | Task 1 0.7015 | 112,057 proteins | higher = more disordered | 05_benchmark/benchmark_spec_v2.json |

^a^ Task 1 labels AlphaFold mean pLDDT < 70; AlphaFold is the reference, not ground truth.

^b^ Task 2 uses the distributed ddG unchanged: dG(mutant) - dG(wild type), positive = stabilising.

^c^ Leave-one-species-out covers 19 species: rye (2 labelled proteins) and cowpea (19) were in training and are skipped only here.

^d^ SD is ddof = 1 for the leave-one-species-out row and ddof = 0 for the cross-validation row, following the value each was frozen at.

^e^ Task 2 is scored on a frozen 448-domain split and evaluates the released model. The validation row against measured folding free energies scores substitution TYPES, not the model, over 926 domains. The two figures, +0.4438 and 0.1227, are therefore different quantities on different data and are not comparable.

*Built by VERIFICATION_SCRIPTS/tables/TS8_validation_benchmark.py.*

**Table S9** Comparison of CropStabDB with related resources

Comparison of CropStabDB with related resources. Each row states what that resource reports, how far it reaches, whether it needs a three-dimensional structure, whether it names a residue to change, and whether it covers plant proteomes. Values for other resources are as reported by their own publications; the CropStabDB row is recomputed from the release database.

| **Resource** | **What it reports** | **Coverage** | **Needs a structure** | **Names a residue** | **Plant proteomes** | **Ref** |
| --- | --- | --- | --- | --- | --- | --- |
| Meltome Atlas | measured melting temperature | 48,000 proteins, 13 species | No | No | No | [1] |
| Mega-scale proteolysis | measured folding free energies | ~776,000 variants of 479 domains, 40-72 aa | No | Yes | No | [13] |
| ProTstab2, DeepTM, DeepSTABp | predicted melting temperature | proteome scale | No | No | No | [9-11] |
| FoldX | dG change of a supplied substitution | one protein at a time | Yes | supplied by the user | No | [27] |
| AlphaFold DB | per-residue model confidence | proteomes across the tree of life | produces them | No | Yes | [12] |
| Protein language models | predicted structure from sequence | >617 million sequences | No | No | not specifically | [28] |
| Gene shuffling and directed evolution | measured variants | one protein | No | screened, not named | Yes | [5,6] |
| Manual orthologue alignment | designed substitutions | one protein | No | Yes | Yes | [7] |
| Ensembl Plants | sequences and annotation | plant proteomes | No | No | Yes | [14] |
| CropStabDB | folding propensity in six interpretable components, plus orthologue-derived candidate substitutions | 1,425,716 proteins, 30 species, 12 families; 2,097,576 candidates in 51,440 proteins | No | Yes | Yes | this work |

^a^ "Names a residue" asks whether the resource identifies which position to change. FoldX evaluates a substitution the user has already chosen; directed evolution screens variants rather than naming one.

^b^ The AlphaFold Protein Structure Database does not require a structure because it produces them; CropStabDB stores its per-residue confidence as a reference layer.

^c^ Reference numbers are those of the main reference list.

*Built by VERIFICATION_SCRIPTS/tables/TS9_resource_comparison.py.*
