## Supplementary figures and images for "CropStabDB: a protein stability resource and Engineering Atlas of orthologue-supported substitutions for 30 plant proteomes"

### Supplementary_Figure_S1.tiff

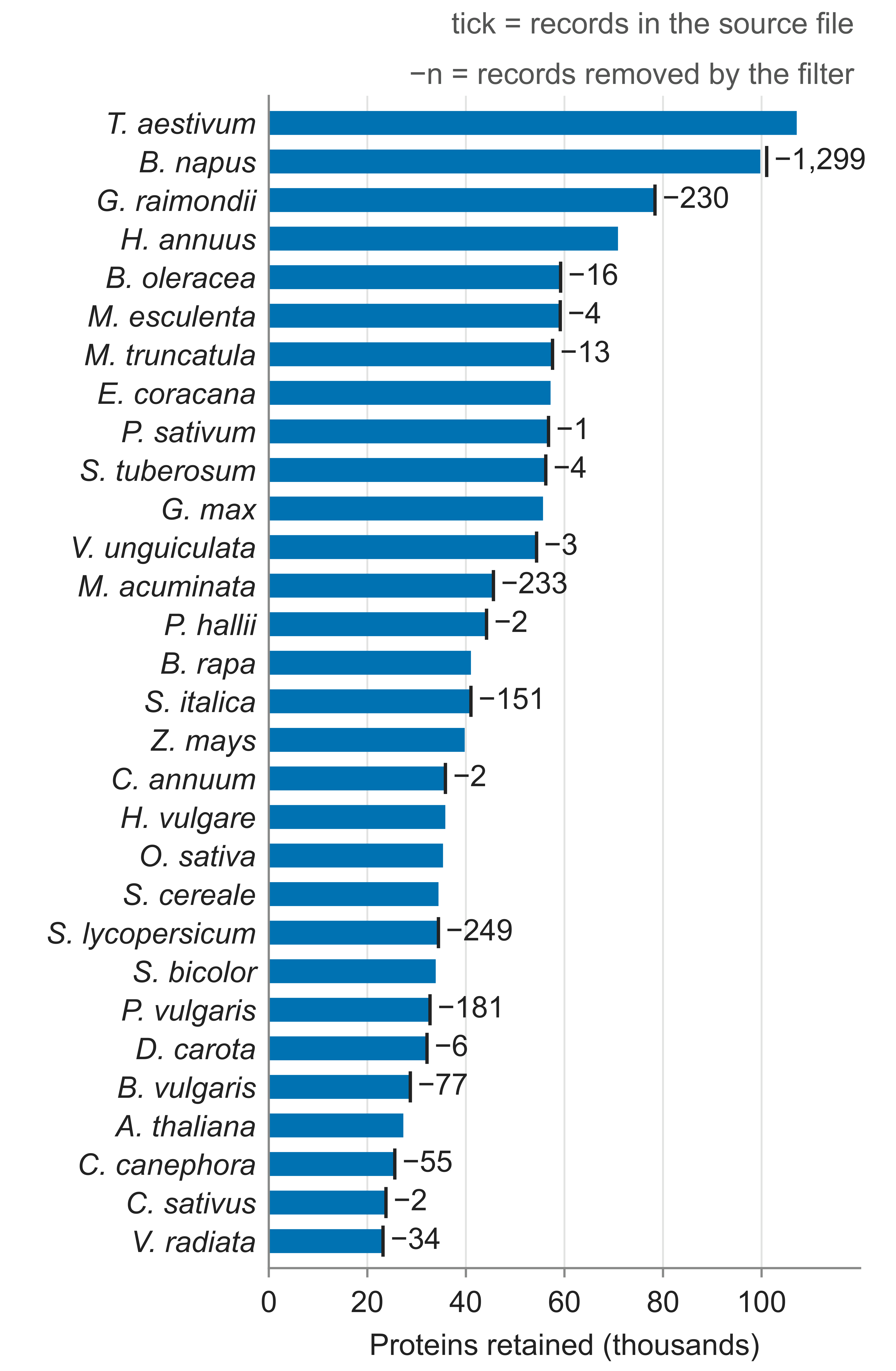

### Supplementary_Figure_S2.tiff

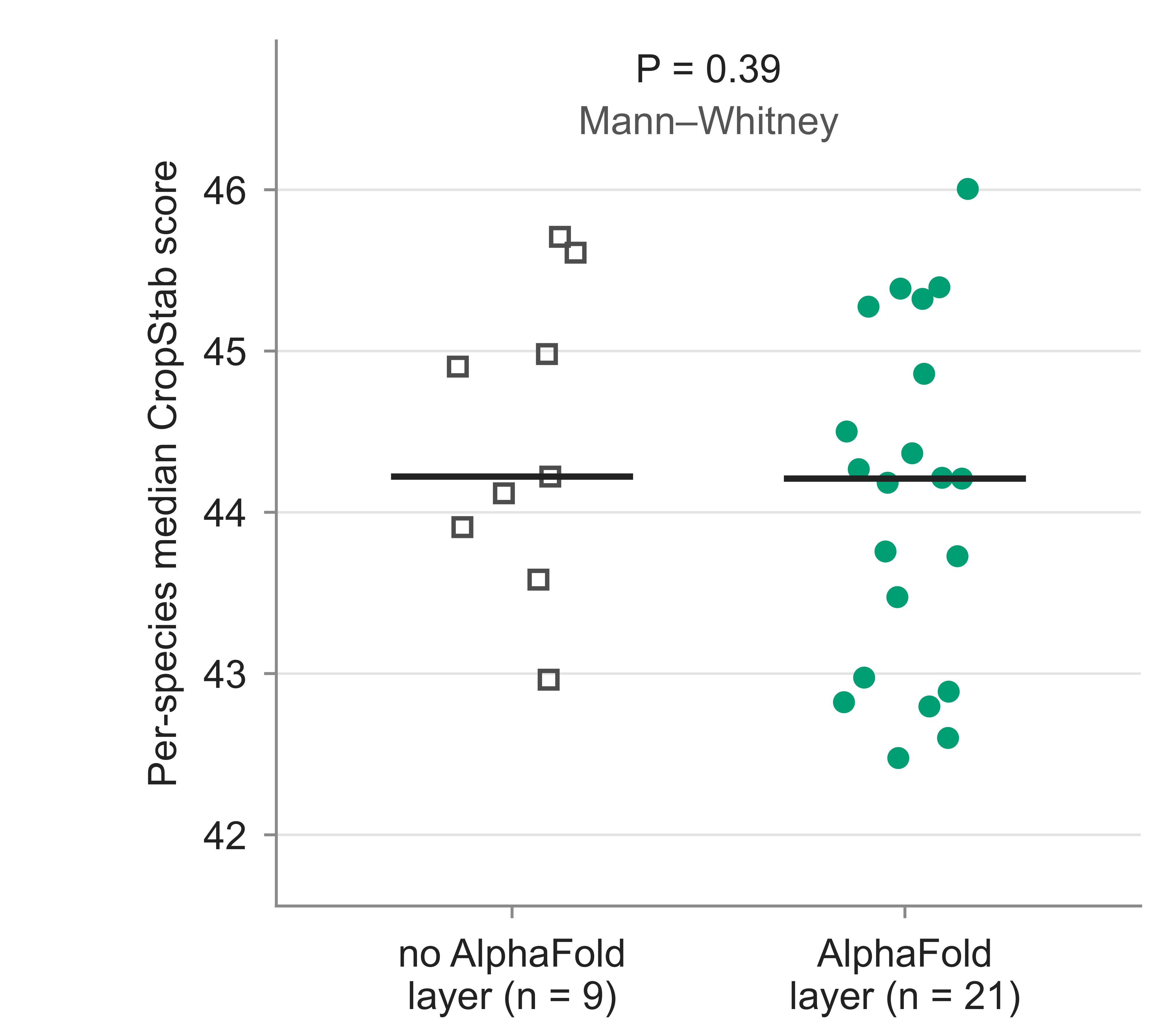

### Supplementary_Figure_S3.tiff

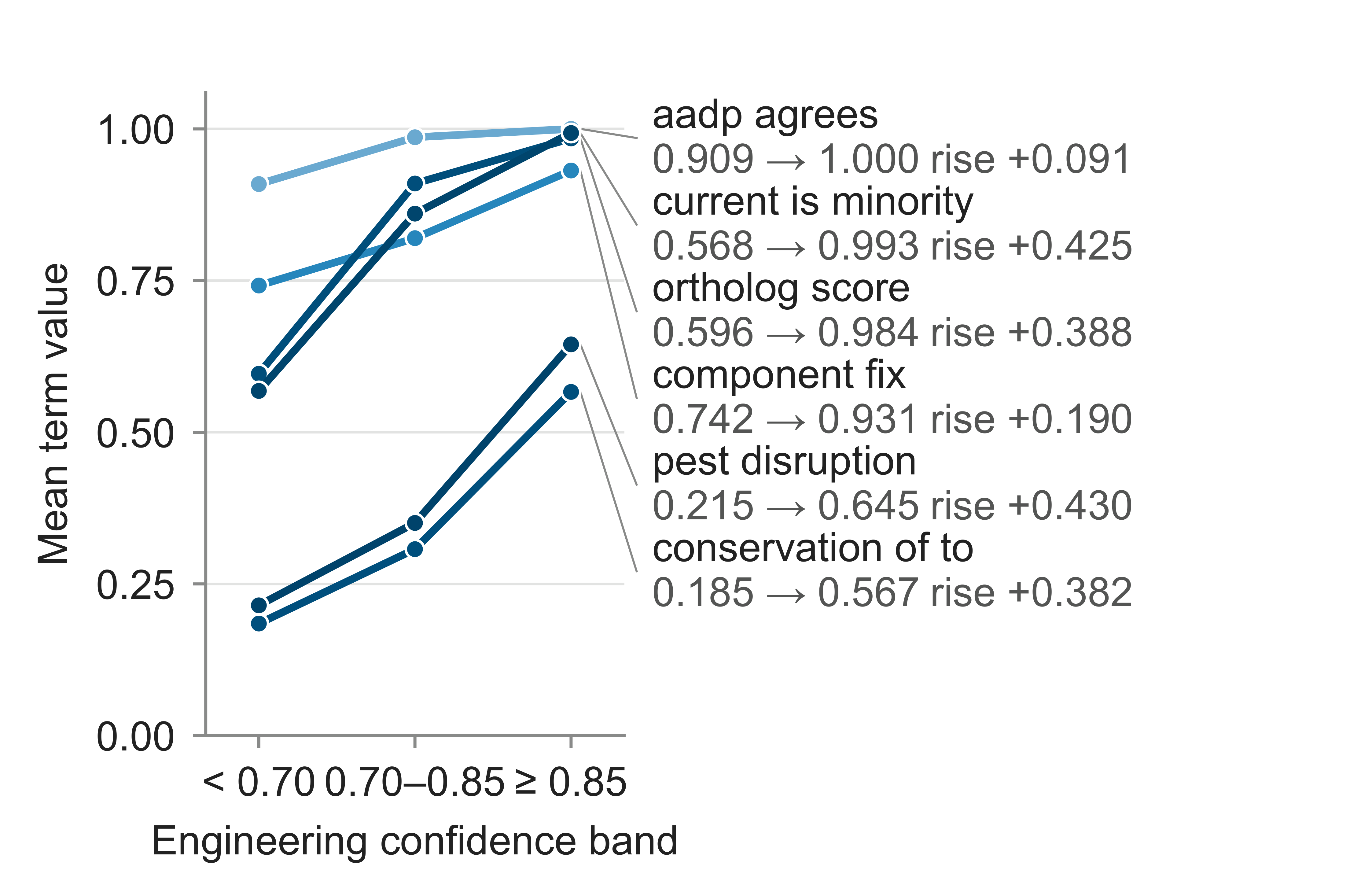

### Supplementary_Figure_S4.tiff

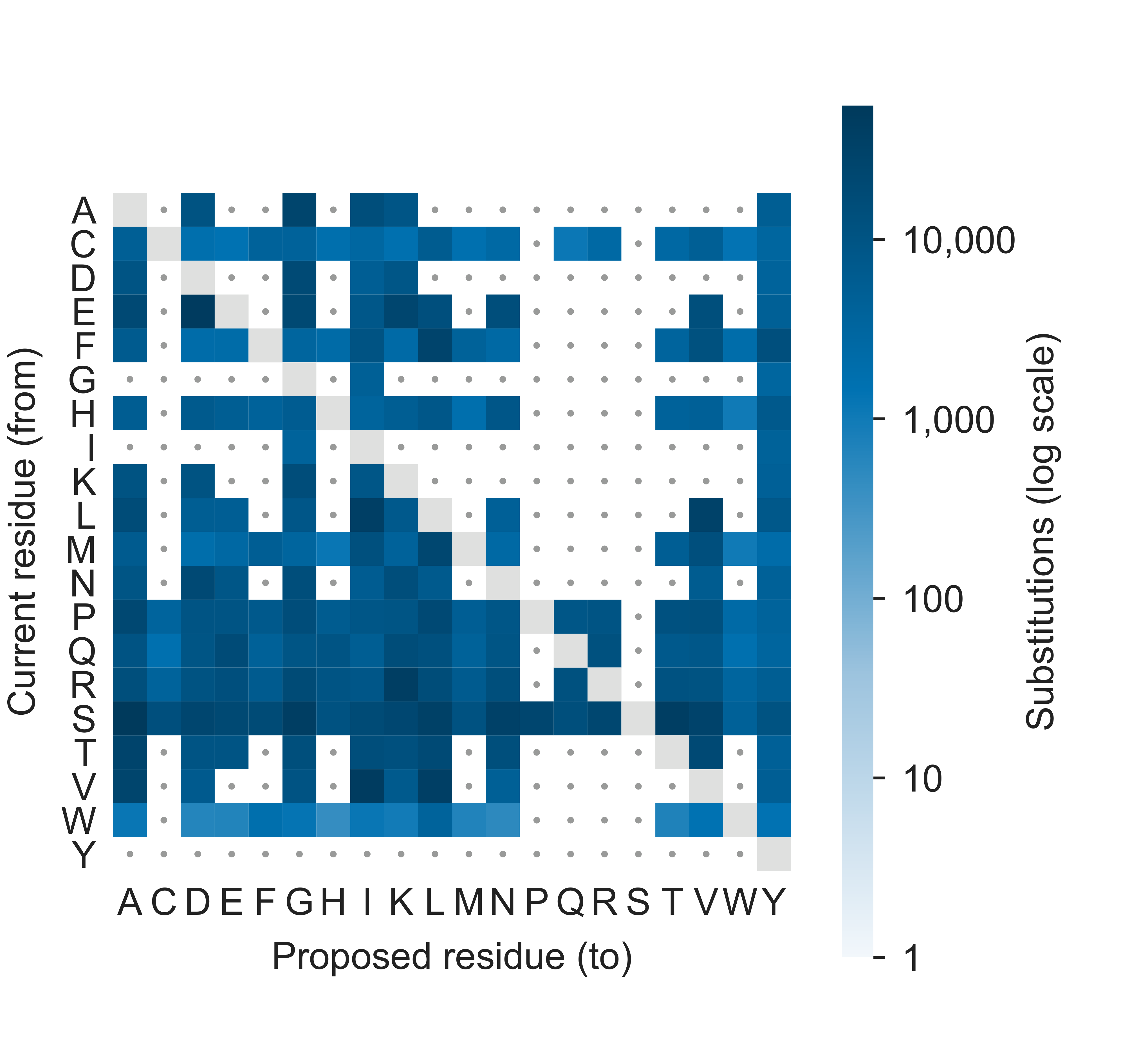

### Supplementary_Figure_S5.tiff

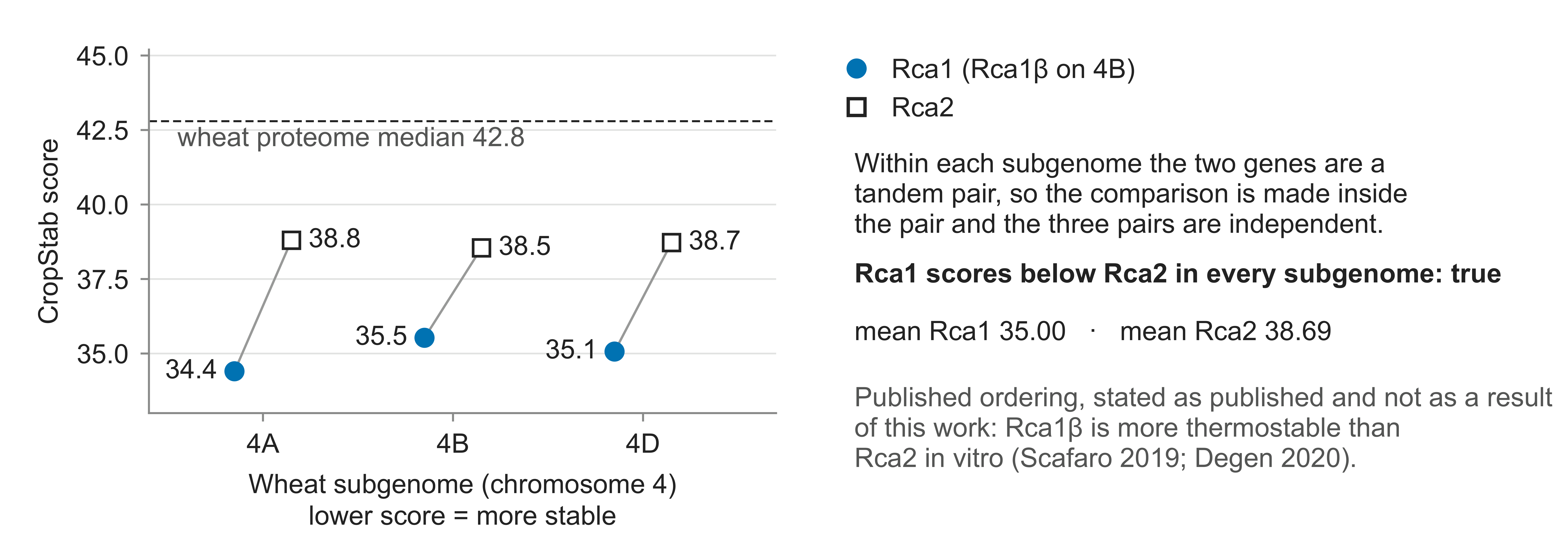
